# A normative account of human temporal structure learning

**DOI:** 10.64898/2026.01.10.698785

**Authors:** Niloufar Razmi, Xufeng Caesar Dai, Leah Bakst, Matthew R. Nassar

## Abstract

People rapidly recalibrate their expectations about the world in the face of surprising observations. This recalibration should depend on the temporal structure of the environment, however how people should and do learn temporal structures remains unknown. To examine this gap, we developed a Bayesian model that infers temporal structure of the environment directly from observations and makes accurate predictions across qualitatively different environments. We tested predictions of our model in an online behavioral study in which participants predicted outcomes generated according to various temporal structures, and demonstrated that people learned to exploit these structures in a qualitatively similar manner to that of our structure learning model. Furthermore, we show that normative structure learning accounts for puzzling asymmetries previously observed in sequential effects of temporal structures on human learning. Taken together our model and empirical data provide the first general account of how people use temporal structures to interpret unexpected observations in the service of learning.

## Introduction

Successful decision making requires forming accurate predictions about important environmental variables, such as potential gains or losses associated with an available action (Behrens et al., 2007; Pulcu & Browning, 2017). Doing so is challenging because the environment often changes and the structure of such changes is typically unknown. For example, when food ordered from one’s favorite restaurant doesn’t match the expectation, the bad experience might be attributed to a one-off event, a longer lasting effect that might at some point in the future revert back to normal, or a permanent change in the restaurant’s quality (Fig.1). In each of these scenarios, the environment undergoes change, but each has its own distinct temporal structure such that unexpected observations have different implications for the future in each scenario, and thus demand different learning policies. While normative approaches have revealed how learning should be modulated in environments that change under specific structural assumptions (Behrens et al., 2007; Nassar et al., 2010; O’reilly, 2013; Piray & Daw, 2020), it remains unknown how these structural assumptions could, themselves be learned, or whether people are able to do so in order to optimize their predictions and related behavior.

**Figure 1.**
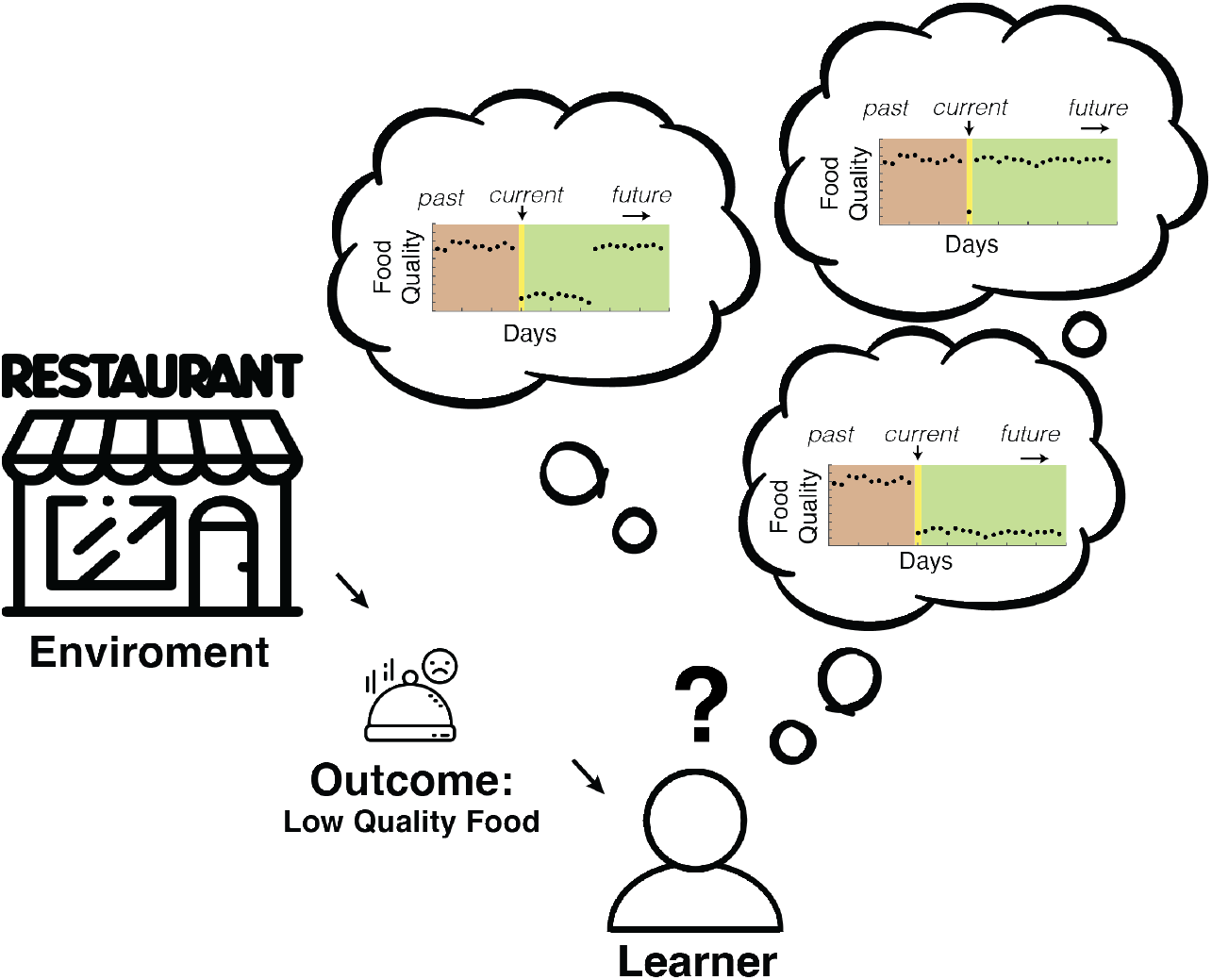
Structural knowledge of an environment is important for optimal learning: In the real world, people are constantly faced with the problem of predicting future outcomes from past experiences. But how to best update such predictions after a new experience depends critically on assumptions about the underlying structure of the environment. For example, here, a person is trying to predict the quality of the food from a given restaurant based on observations from previous days. The same unexpected outcome (low quality food) can be interpreted in at least three different ways according to differing structural assumptions: the bad food will persist indefinitely (perhaps there is a new management who has decided to cut back on expenses by using cheap products), the bad food will persist for a while but will go back to normal after a certain period (perhaps there are two head chefs working with alternating schedules) or the quality of the food will immediately go back to what it was on previous days (there was a malfunctioning oven in the kitchen today that will be fixed tomorrow). This creates a type of “meta-learning” challenge: how do we learn the temporal structure of an environment so as to know which assumptions allow for the best predictions of future observations?

Bayesian models have proven useful in providing normative prescriptions for learning under uncertainty, but have typically embedded fixed assumptions of the test environment that limit their generalizability (Behrens et al., 2007; A. Collins & Koechlin, 2012; Konovalov & Krajbich, 2018; McGuire et al., 2014; Nassar et al., 2010; Piray & Daw, 2020). For example, Bayesian models have provided prescriptions for optimal learning in environments with volatility and stochasticity (Behrens et al., 2007; Piray & Daw, 2021), changepoints (Adams & MacKay, 2007; Nassar et al., 2010; Wilson et al., 2010), reversals (A. Collins & Koechlin, 2012), oddball events (d’Acremont & Bossaerts, 2016; Nassar et al., 2019) and sequences (Konovalov & Krajbich, 2018). These models differ from one another in their assumptions about the sorts of transitions that are expected to occur in the environment, for example in terms of whether abrupt state transitions are expected to occur, whether they will persist in time, and whether they will return to previously encountered state. Thus, the models account for aspects of normative learning behavior that are only observed (and indeed only normative) in certain environments, leaving open the question of whether and how people learn which structural assumptions to rely on in real world scenarios where the environmental structure must be learned directly from the observations.

In principle, Bayesian inference could be used to solve the more general structure learning problem if it were applied to a generative framework sufficiently flexible to give rise to the different structural motifs that people encounter in the real world. Indeed, recent work has employed Bayesian models with increasingly flexible assumptions about the environmental structure, however to date such models have been employed using a fixed set of hyperparameters, chosen either to match the task at hand or the behavior of an individual participant (A. Collins & Koechlin, 2012; Heald et al., 2020). The structure learning problem articulated above could be thought of in this framework as the problem of flexibly learning the hyperparameters of a generative model from the observed outcomes themselves.

Here, we develop a normative solution to structure learning through this approach and apply it to a testbed of predictive inference tasks that are particularly well suited for characterization of human learning behaviors, allowing us to test the key behavioral predictions of normative structure learning. We first formulate “temporal structure” in terms of the transition matrix over the “latent states” generating trial outcomes. We use the term latent state, a categorical variable in the generative process, since in the real world, its identity is usually hidden from the learning agent and must be inferred from noisy observations (Gershman & Niv, 2010; Wilson et al., 2014). We then apply Bayesian inference over a highly configurable generative process capable of producing transition matrices characteristic of different environments under different parameterizations (Heald et al., 2020). By doing inference over these parameters, our model effectively learns inductive biases over possible transition matrices that enable it to better infer which types of transitions are likely to occur, even for transitions it has never observed (i.e. the first time entering a new state).

We tested the performance of this model in four predictive inference tasks that differed qualitatively in their generative structure (changepoints, oddballs, reversals, sequences) as well as their demands for adaptive learning. Here, we introduced a unified definition of temporal structure by using different transition matrices to *generate* outcomes for all tasks. We then trained our normative Bayesian model to *infer* the parameters of its generative process that can be used as a prior for any transition matrix. The model achieved high levels of performance across structures and developed different hallmarks of efficient learning for each. We tested behavioral predictions of the model in humans who performed uninstructed versions of the same tasks and found that human behavior qualitatively matched that of our normative model, but deviated quantitatively in part due to heterogeneity in the degree of environmental adaptation across individual subjects. Performance improvements in both the model and people depended on the structure being learned, with initial learning behaviors most closely matched to the demands required for the changepoint structure, and thus greater performance improvements for the other structures. Furthermore, pre-training the normative model on one structure affected its ability to learn another, and these pretraining effects allowed the model to account for puzzling asymmetric sequential effects previously observed in the implicit learning of sequentially presented changepoint and oddball structures (Bakst & McGuire, 2023).Taken together, our results provide a normative account of how people can and do learn the temporal structure of the environment and exploit it to improve their predictions about the world. This work thus bridges the extensive research that has examined adaptive learning in fixed structures in laboratory settings to the real-world scenario in which structure must be learned through repeated experience.

## Results

Here we examine how people should and do learn about the temporal structure of their environment in order to make predictions within it. Our results are organized as follows. First, we describe a normative model of structure learning and evaluate its performance on a testbed of predictive inference tasks. Next, we examine behavioral hallmarks of structure learning that emerge in the model, and compare them to those of human participants performing uninstructed versions of the same tasks. Finally, we examine how structural inferences of the model are affected by pretraining on an alternate structure, and show how such pretraining effects provide a normative explanation for counterintuitive learning asymmetries previously observed in human experiments involving multiple structures encountered sequentially.

### A unified framework for studying temporal structures

We examine the question of how one “should” learn from observed outcomes without a-priori knowledge of their underlying generative structure. To do so, we view learning through the lens of predictive inference, where the goal of learning is to use a new observation to better tune expectations for future ones. Within this framework we formalize the intuitive notion of different “types” of dynamic environments in terms of different classes of transition matrices that might describe the environment. In particular, we consider a mathematical framework that assumes observations *x*_*i*_ are generated from a latent state identified by a categorical variable that can change according to a transition matrix. Learning the full transition matrix would thus enable the model to evaluate the likelihood that surprising observations reflect specific state transitions. Beyond this, learning higher order statistics of the transition matrix might allow for efficient interpretation of new observations even when they are completely inconsistent with previously observed states – for example by determining whether an outlying data point is more likely to be a lasting fundamental change in the environment (changepoint) or a transient shift unlikely to persist (oddball). In short, our framework formalizes the problem of learning temporal structure in terms of learning the transition probability matrix over a set of latent states.

To assess learning we extend an established predictive inference task framework (Fromm et al., 2023; McGuire et al., 2014; Nassar et al., 2010, 2021) (Fig.2a) to study four different temporal structures (Fig.2b-e). The task requires participants to move a bucket to the most likely horizontal location of a helicopter, which is occluded by clouds, but which drops a bag containing gold coins on each trial. Our extensions of the task included cases in which helicopter movements were governed by changepoint (Fig.2b), oddball (Fig.2c), reversal (Fig.2d), or sequence (Fig.2e) generative structures. These four structures are frequently studied individually in psychology tasks and each structure has characteristics that make unique demands for how an agent should adjust its learning in response to an unexpected observation.

**Figure 2.**
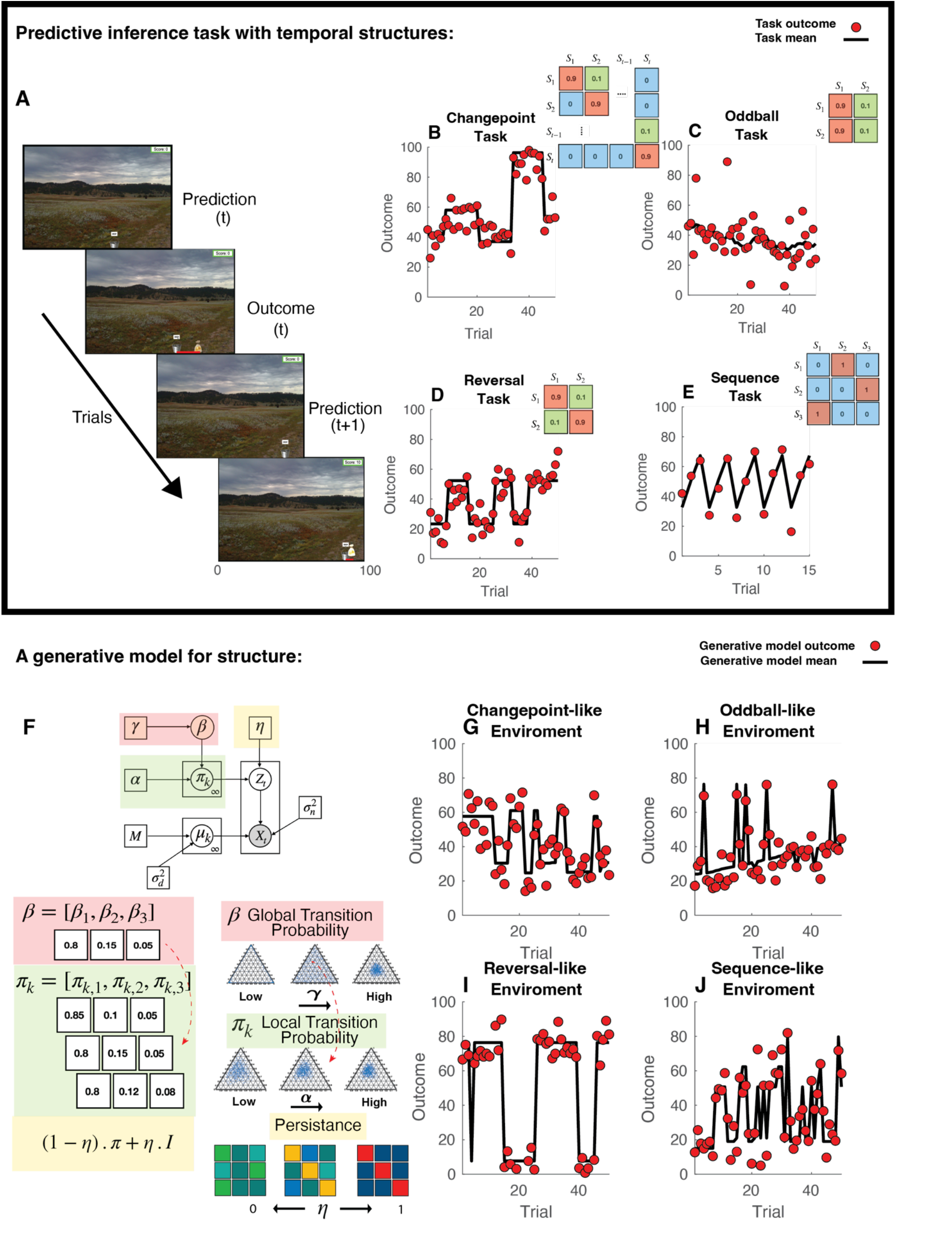
Temporal structure learning as inferring the hyperparameters of a generative process for transition matrices: A) Screenshots of a predictive inference task, framed as inference about the location of a helicopter dropping bags of gold from the sky. On each trial, participants report their belief about the helicopter position by moving a bucket to their inferred location (prediction). Participants then observe the actual horizontal bag location (outcome) which is drawn from a distribution centered on the helicopter. B-E: The helicopter and bag positions change over time according to one of four different generative structures. Helicopter (black) and bag (red) positions (ordinate) are plotted across trials (abscissa) for example changepoint (B), oddball (C), reversal (D), and sequence (E) tasks. Helicopter transitions for each task can be described by a transition matrix displayed inset. **F:** A persistent hierarchical Dirichlet process (pHDP) provides a flexible generative architecture for creating transition matrices and task data (top). The pHDP generates transition matrices that depend critically on set of hyperparameters *γ, α, η, σ*_*n*_, *σ*_*d*_ where *γ* and *α* are the concentration parameters of the two Dirichlet Processes (DPs), *η* is a persistence factor and *σ*_*n*_ and *σ*_*d*_ are standard deviations of noise and drift respectively. The pHDP draws a vector (*β*) of probabilities from the high-level DP parameterized by *γ*, such that higher values of *γ* result in more states with a reasonable associated probability (red, also see bottom left panel). *β* serves as the base of a second DP with parameter *α*, and drawing from this DP yields rows of the transition matrix, where higher *α* would results in rows more similar to original β (green, bottom left panel). On each trial, the pHDP flips a coin with weight *η* determining whether states will transition according to π or whether the current state will persist, such that the resulting transition matrix reflects a weighted mixture between π and the identity matrix *I*. (**G-J**) State means (black) and outcomes (red) simulated from the pHDP using four different sets of hyperparameters that led to outcomes resembling those from the four tasks shown in B-E.

In the changepoint condition, the environment undergoes abrupt transitions to new, non-recurring states. As such, extreme prediction errors are often indicative of persisting changes that demand rapid learning (i.e. higher learning rate), whereas more modest prediction errors are indicative of random noise demanding slower learning(Nassar et al., 2010). In contrast, in the oddball task (Fig.2c) an extreme prediction error does not warrant the same response, instead incentivizing learners to de-emphasize outlying datapoints, as such oddball events are not predictive of future observations (d’Acremont & Bossaerts, 2016; Nassar et al., 2019). Reversal tasks (Fig.2d) and sequence tasks (Fig.2e) both demand reuse of information from previously visited states, but differ in the degree to which a given state persists in time before transitioning to a new state.

These qualitative differences in the temporal structure of the four environments can be formalized through their *transition matrices*. A transition matrix is a |S| x |S| matrix, with |S| being the number of states, where each row corresponds to the probability of transitioning from a state to any other state and therefore the sum of each row of the matrix equals 1. In our task, the transition matrix describes transitions from one helicopter position to another, and since the helicopter is not directly observable, the transition matrix is over latent, rather than observable, states. In the changepoint task, the transition matrix includes a high probability of staying in the current state with a low probability of transitioning to a completely new state (Fig.2b, upper right). In the oddball task, there are two states and transitions to one of the states are always more probable regardless of the current state (Fig.2c, upper right). In the reversal task, there are two states, with a high probability of self-transitions in each state, and a lower probability of transitioning between states (Fig.2d, upper right). Finally, in sequence learning task, the transition matrix is deterministic, such that each state transitions to the next one with the probability of one (Fig.2e, upper right). Thus, the latent state transition matrix enables us to describe four common psychology task structures using a single unified mathematical framework, and thereby define the structure learning problem in terms of learning the characteristics of a transition matrix.

To model structure learning with Bayesian inference, we took a two-step approach. First, we defined a compact generative process capable of expressing a wide range of transition matrices through modulation of a small number of hyperparameters. Second, we used Bayes rule to invert that generative process allowing us to infer the latent states, transition matrices, and hyperparameters simultaneously.

The persistent Hierarchical Dirichlet Process (pHDP) provides a compact framework capable of generating a wide range of transition matrices (Fig.2f top). The pHDP is hierarchical in the sense that it includes a high and low level Dirichlet process (DP), each of which generates a set of probabilities controlled by its own single concentration parameter. In the high level DP the concentration parameter γ controls the degree to which state occupancy probability β is concentrated within a small number of states (Fig.2f left red panel). The low-level DP samples a probability vector centered on the overall state occupancies β for each state corresponding to the transition probabilities out of that state, and α controls the concentration of that sampling distribution. Collectively, these vectors give rise to a transition matrix π, (Fig.2f left, green panel) with α controlling the similarity between columns in that matrix. Finally, a persistence parameter, *η*, controls an additional process that can give rise to self-transitions (Fig.2f, right yellow panel). This corresponds to taking a weighted average of the transition matrix π and an identity matrix (Fig.2f left yellow panel). While this transition matrix defines a distribution over the categorical states in the task, each latent state has a unique mean μ (sampled uniformly from 𝒰[0,100] for each categorical state, with a trial-wise drift rate of σ_*d*_), and outcomes are generated from a Gaussian distribution 𝒩(μ, σ_n_). The pHDP model is extremely expressive considering its small number of hyperparameters and can generate transition matrices and outcomes resembling the changepoint, oddball, reversal and sequence tasks described above using different hyperparameter settings (Fig.2g-j).

### Inference on the pHDP enables temporal structure to be learned through experience

Given that outcomes generated from a pHDP under different hyperparamter settings cover a wide range of task structures that differ in their demands for adaptive learning, we next examined whether Bayesian inference over the pHDP could infer structure and implement structure-specific adaptive learning. We implemented an online Bayesian inference model that updated a posterior distribution over hyperparameters, the transition matrix, the current latent state, and mean outcomes associated with each latent state simultaneously. The inference model started with a uniform prior over its hyperparameters and updated this distribution with each new observation in order to build expectations about the sorts of transitions that should be expected in a given environment. While exact inference over the generative structure is computationally infeasible, we implemented it using a set of approximations, including taking a particle filtering approach to the problem of representing exact state sequences. The particles were propagated, weighted according to their likelihood of giving rise to the newly observed outcome, and normalized at the end of each trial. We refer to this inference model as pHDPi to distinguish it from the generative model itself.

Predictions made by the pHDPi on example tasks suggest that it learned the appropriate temporal structure within a single task session. To test whether the pHDPi could infer and exploit temporal structures, we generated tasks that included either changepoint, oddball, reversal, or sequence structures and allowed the pHDPi to perform each of these tasks by observing one outcome at a time and predicting the next one. Results from an example set of these tasks reveal that model predictions change considerably over the course of a single learning session (compare Fig.3a-d to Fig.3e-h). Maximum likelihood predictions of the mean (black line) and posterior predictive distributions (heatmap) diverged considerably from outcomes (red points) early in the task (Fig.3a-d), but came to match them by the end of the simulated 200 trial example sessions (Fig.3e-h). The pHDPi started with the same qualitative predictive behaviors for each task, but gradually with experience, developed differentiated behaviors across the different tasks. These differences were most notable in response to changes in the latent state, which are identifiable through the large shifts in the outcomes that accompany them.

**Figure 3.**
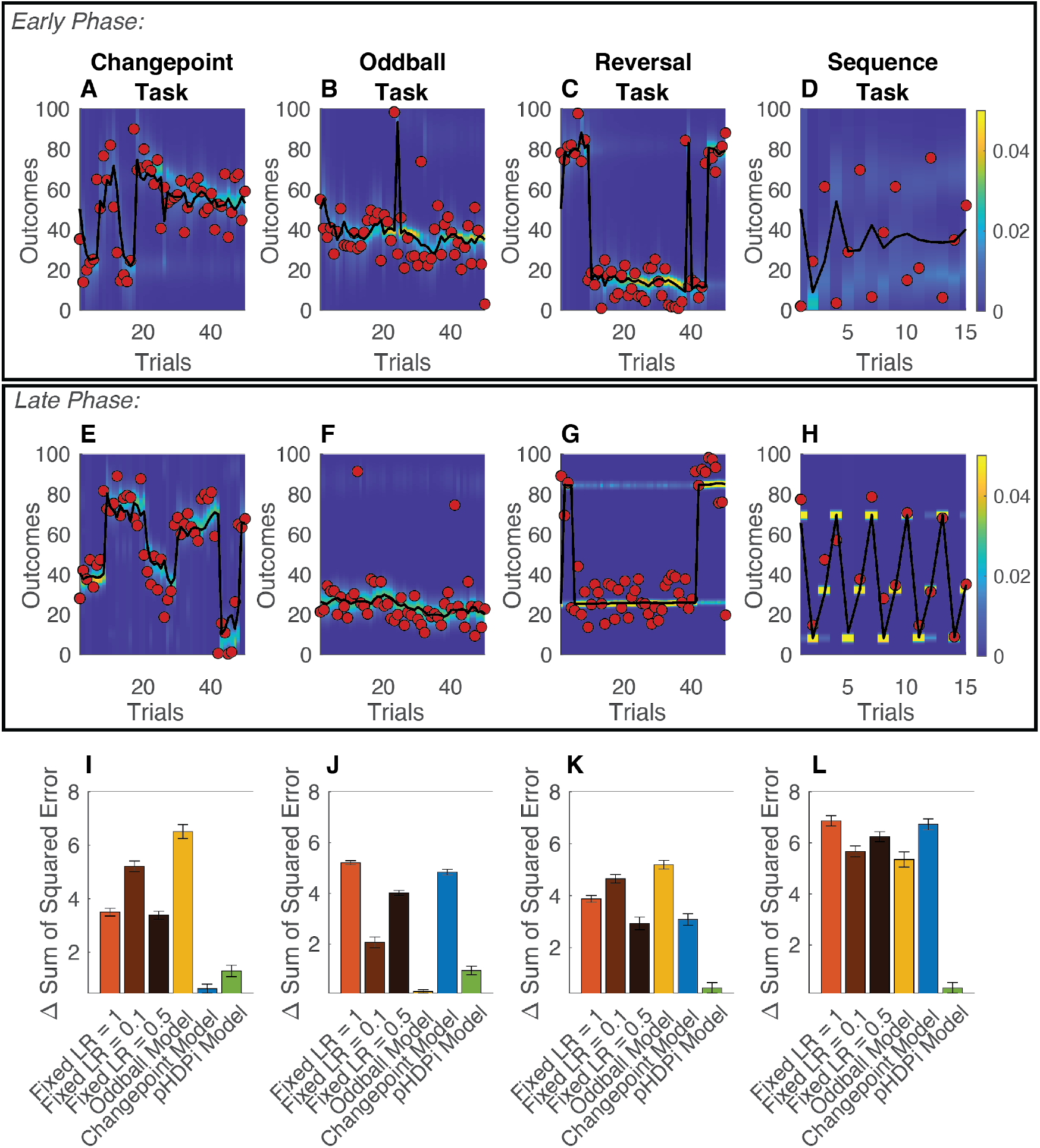
The persistent hierarchal Dirichlet process (pHDPi) model learns the four tasks: Example of responses of the model trained in the four different tasks separately, divided by early phase of the task (the first 50 trials, in sequence task only 15 trial shown for better visibility) in A-D and late phase of task (the last 50 trials, in sequence task only 15 trial shown for better visibility) in E-H. Red dots denote the outcomes, the heatmap represent the posterior probability of model’s belief over the mean of the trial and black line shows the expected value of the mean. We can see qualitative differences in the behavior of the model in late phase compared to the early phase of the experiments consistent with how we expect an ideal observer would respond in each respective task structure. (I-L) Comparison of relative sum of squared error (where the minimum SSE in each run was subtracted from all models’ SSE) of the model to some preexisting models: three delta rule model with a fixed learning rate of 0.1, 0.5 and 1 and Bayesian models designed for the changepoint and oddball tasks.

The pHDPi model performed better than simple and structure-specific models when compared across task structures. When compared with delta rule models that update predictions as a fixed proportion of prediction errors on each trial the pHDPi model performed better across all four tasks (3I-L). The pHDPi also compared favorably to structure specific learning algorithms, specifically Bayesian models that were designed for the change-point and oddball tasks. These algorithms performed well in the environments for which they were designed but failed to generalize to other tasks with different temporal structures (Fig.3i-l blue and yellow bars). Thus, the pHDPi achieved robust predictive accuracy across our distinct environments that differed in their demands.

Having shown that the pHDPi flexibly learned our task set, we next examined how it did so and found that the model inferred different hyperparameters for each task, thereby giving it different expectations about how the transition matrix would change in each. Qualitatively, learning led parameters to diverge from those in the initialized model, but the direction of divergence differed across tasks (Fig.4a). Parameters for some tasks diverged more than others, with the changepoint task notable in the similarity of its posterior parameters to the prior, as measured by Euclidean distance (Fig.4b; median distance in changepoint task: 6.12, SEM = 0.37, median distance over other tasks: 7.21, SEM = 0.17, p=0.01). One interpretation of this result is that the changepoint task required less hyperparameter learning than the three other structures examined.

**Figure 4.**
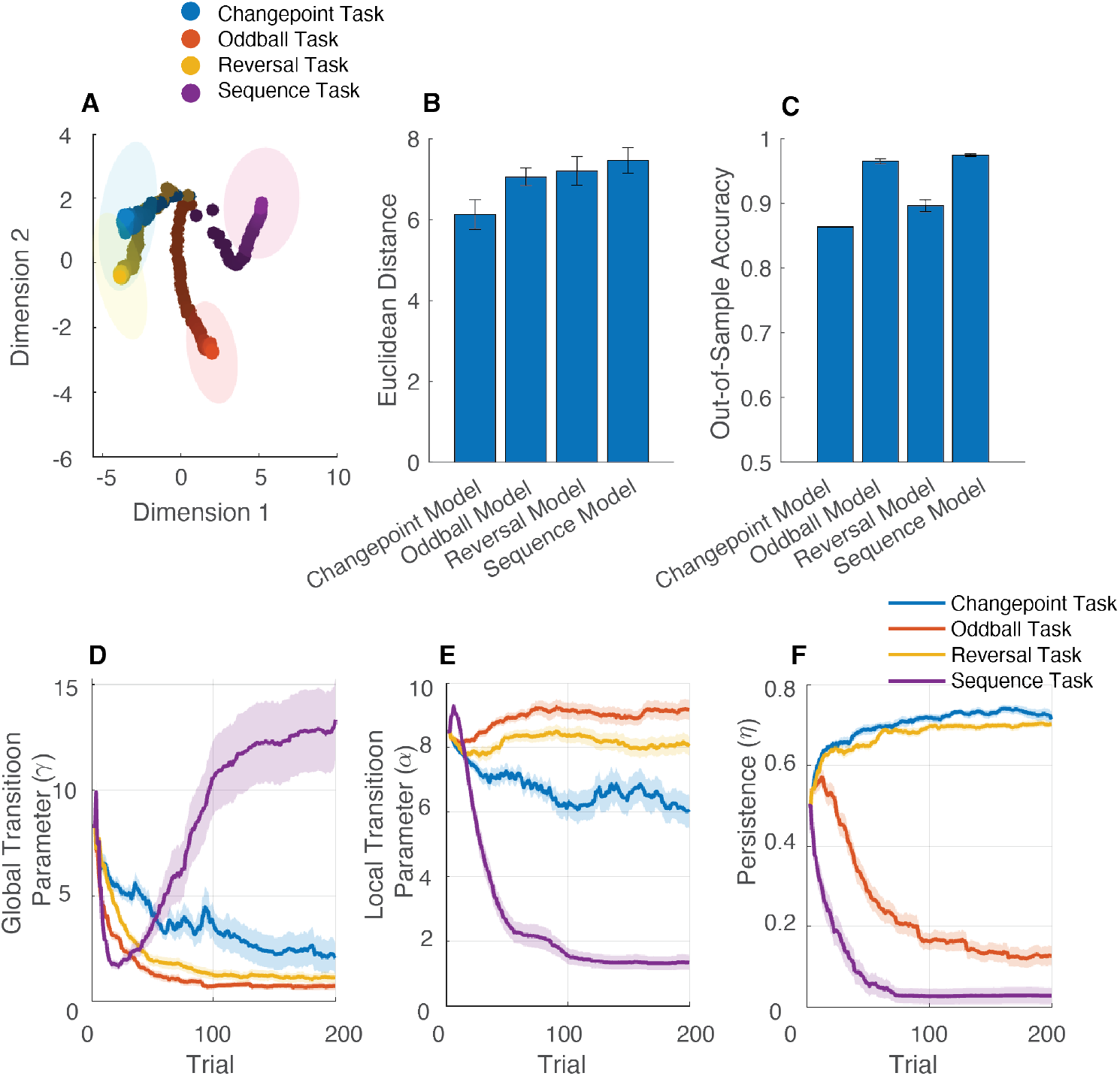
Inferred hyperparameter distributions differentiate the four tasks to different degrees. (A) Multidimensional scaling (MDS) was used to visualize the expected value of the model’s hyperparameters (averaged across 40 runs of each task) by transforming the Euclidean distance between pairs of tasks’ 5 hyperparameters to distance between task pairs in two MDS dimensions. The hyperparameters are z-scored and the initial values of each hyperparameter are subtracted before performing the MDS. The resulting two dimensions of the MDS are also in units of standard deviation. The color hues correspond to different tasks whereas the shades correspond to trials within an experiment. Note how all tasks start at the same point in the 2 dimensions (darker shade points) but gradually separate from one another (lighter shade points). The confidence ellipses represent the covariance of the 40 different runs of each task for only the last trial in the experiment. (B) We quantify the original average Euclidean distance in 5 dimensions (bars) with SEM (error bars) for the model trained each task. Here, the average distance from starting point is significantly lower in the changepoint task compared to all the other tasks (an observation also qualitatively evident in Fig.4a). (C) shows the 10-fold cross-validation accuracy of a SVM binary classifier trained separately on final learned hyperparameters to classify each task vs all the other tasks (with bars showing average accuracy across the 10 folds and error bars depicting the SEM). In D through F, we plotted the three hyperparameters related to the transition matrix (*γ, α, η*) averaged across 5 runs of each task with shaded region showing SEM. Different values of the three main hyperparamters of the pHDP model correspond to different task structures.

To quantify the separability of the learned hyperparameters, we trained a support vector machine to perform binary classifications for each task structure (in comparison to all others) using the final timestep hyperparameters learned in that task structure (Fig.4c). Classification accuracy was relatively high for all four task structures, but the sequence and oddball tasks achieved higher 10-fold cross-validation accuracy (mean = 0.97, SEM = 0.0003 and mean = 0.96, SEM = 0.0005 for respectively) than the reversal learning task (mean = 0.89, SEM = 0.0013) and changepoint task (mean = 0.86, SEM = 0.0016). Further investigation of classification errors revealed that confusions were most prominent between the changepoint and reversal tasks, which led to distinct but slightly overlapping learned hyperparameter distributions. Taken together, these results suggest that the pHDPi learned qualitatively different tasks by gradually adjusting its hyperparameters, albeit to a degree and in a direction that depended critically on the task structure.

These hyperparameter adjustments also provided some intuitions about how exactly the model interpreted each task to differ from the others (Fig.4d-f). The γ hyperparameter controls the prior probability distribution over states, with smaller values reflecting a stronger concentration of probability density on the most common states, and larger values stipulating a flatter and wider distribution over states. Interestingly, γ was inferred to be highest in the sequence task, which only contained three states, but in which the probabilities for those three states were equivalent and uniformly high. Thus, the shape of the probability distribution over states appears to have controlled inference of γ more so than did the overall number of states occupied, at least in the case of the sequence task. Relatively high values of γ were also inferred for the changepoint task, consistent with large number of unique states encountered in this task structure. The α hyperparameter reflects the similarity of rows in the transition matrix, and consistent with this, was inferred to take the lowest values in the sequence task where transitions were completely dependent on the current state (Fig.2e). The *η* persistence hyperparameter was inferred to be high in changepoint and reversal tasks, in which states persisted for multiple timesteps in a row, and low for the sequence task, in which they did not persist (Fig.4f). Inferred values of *η* were also low for the oddball task, likely allowing the model to capture the transient nature of the oddball events, but leaving the frequent repetition of non-oddball trials to be explained by a relatively high concentration of probability associated with the most common states (see γ, Fig.4d). Inspecting the slope of the learning curves for the three hyperparameters shows a sudden change around trial 50, especially in case of oddball and sequence tasks, compared to a more gradual change for changepoint task, suggesting an almost phase-transition phenomenon for oddball and sequence tasks which is consistent with the dramatic change in behavior of the model after learning these two tasks (Fig.4d-f). It is noteworthy that, while all hyperparameters were recoverable to some degree, the γ parameter was recovered less reliably than the others, muddying its interpretation slightly (see Supplementary Fig.1).

Overall, the pHDPi model made accurate predictions across a variety of different structures and did so by adjusting its beliefs about generative hyperparameters in accordance with observed outcomes. These adjustments differed for each task structure, with some structures requiring smaller adjustments than others (changepoint), but all structures driving unique hyperparameter settings that enabled accurate and structure-specific predictions.

### Can people learn the temporal structure of tasks without explicit instructions?

Given that the pHDPi was able to learn to exploit different temporal structures after observing a relatively small number of samples (number of trials = 200), we next tested whether people were able to do the same. We performed an online behavioral study in which 480 human participants were randomly assigned to perform a predictive inference task requiring them to catch bags dropped from a helicopter that matched one of the four temporal structures (Fig.2a; N=120 per task structure). The horizontal location of each bag corresponded to the outcome in our previous modeling section and the location of the helicopter corresponded to the mean of the outcome generative distribution. Importantly, the only instructions were to move their bucket to the location at which the next bag was expected to fall.

People and the pHDPi both improved their performance over the course of the 200 trials of the predictive inference task. Errors were larger in the early phase of the task (first 50 trials) than the late phase (last 50 trials; Fig.5a). This difference was large and positive for both humans and the pHDPi in oddball (Humans: median *ΔSSE*_*z*_ = 0.56, *σ* = 0.12, *p* < 0.0001, Model: median *ΔSSE*_*z*_ = 0.95, *σ* = 0.10, *p* < 0.0001), reversal (Humans: median *ΔSSE*_*z*_ = 0.12, *σ* = 0.0.09, *p* = 0.02, Model: median *ΔSSE*_*z*_ = 0.94, *σ* = 0.09, *p* < 0.0001 and sequence (Humans: median *ΔSSE*_*z*_ = 0.32, *σ* = 0.0.08, *p* < 0.0001, Model: median *ΔSSE*_*z*_ = 1.02, *σ* = 0.09, *p* < 0.0001 tasks. For the changepoint task, the difference in humans was not reliably different from zero (Humans: median *ΔSSE*_*z*_ = 0.01, *σ* = 0.09, *p* = 0.62, Model: median *ΔSSE*_*z*_ = 0.52, *σ* = 0.12, *p* < 0.0001) suggesting that improvements in the changepoint condition over the session were less dramatic than for the other temporal structures. Indeed the difference was significantly smaller for both humans and the model in the changepoint condition compared to all other tasks (Humans: Changepoint task: *median ΔSSE*_*z*_ = 0.01, Other tasks : *median ΔSSE*_*z*_ = 0.21, *p* < 0.0001, Model: Changepoint task median *ΔSSE*_*z*_ = 0.52, *Other tasks*: median *ΔSSE*_*z*_ = 0.97, *p* < 0.0001). This finding was consistent with the model hyperparameter analysis showing that hyperparameters evolved less in the changepoint condition than the other conditions. While qualitatively similar, average performance improvements in humans were smaller than those in the pHDPi (Humans: overall mean *ΔSSE*_*z*_ = 0.36, SEM = 0.0023, Model: overall mean *ΔSSE*_*z*_ = 1.02, SEM = 0.0024, *p* < 0.0001). This superior improvement of model’s performance compared to humans seemed to arise mostly from the heterogeneity in human participants’ final performance (Supplementary Fig.2). Taken together, these results show that people and the pHDPi were able to improve their predictions over time, and do so to a degree that differs across task structures, with changepoint structures yielding the smallest improvements, consistent with the small relative hyperparameter changes observed in the pHDPi for this task structure.

**Figure 5.**
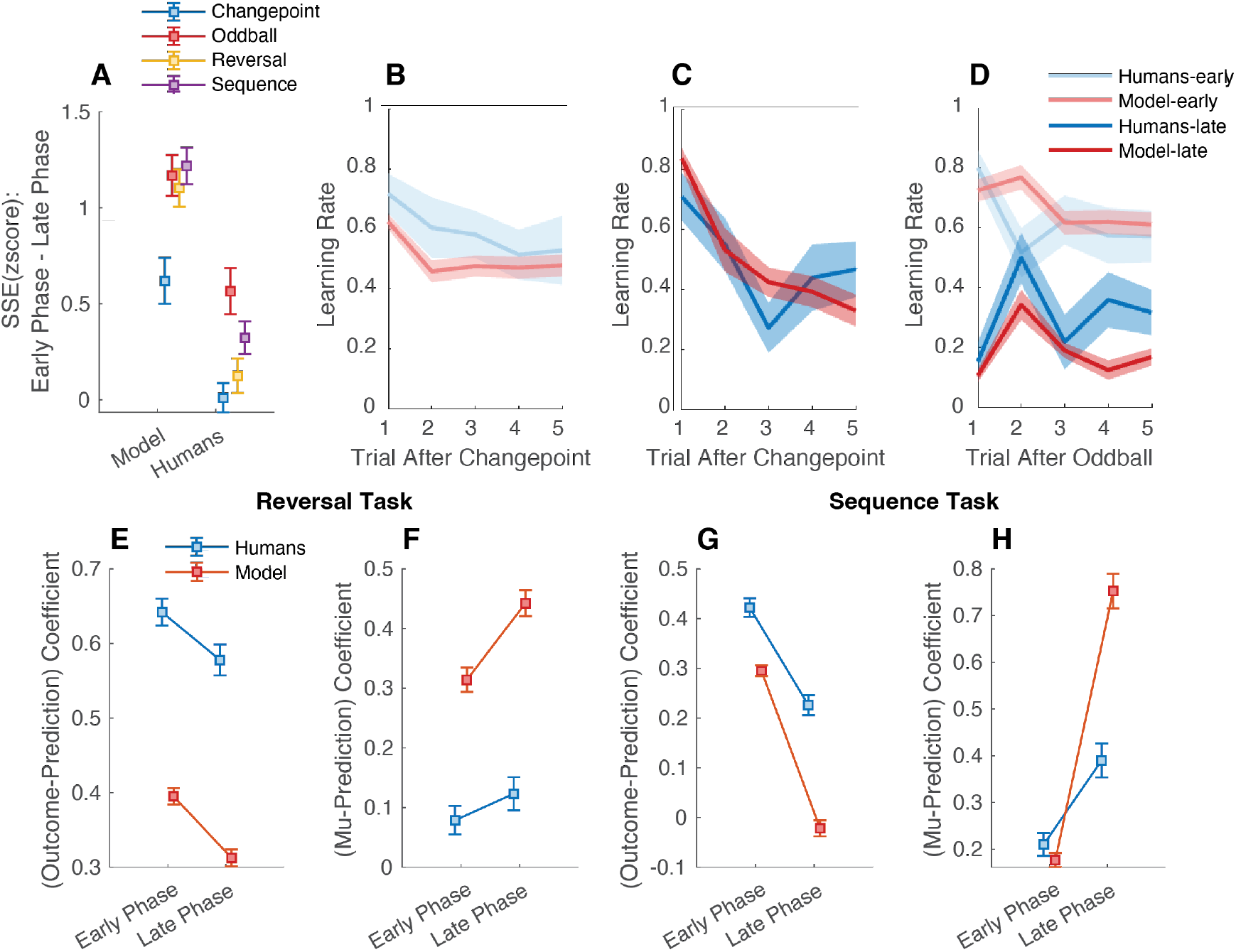
The pHDPi model learns the temporal structure of the tasks quantitatively and qualitatively similar to human participants. A) The differences of (z-scored) sum of squared error (SSE) in the early phase (first 50 trials in the experiment) to late phase (last 50 trials in the task) are significantly greater than zero for all four tasks in model simulations and all tasks except changepoint task in human participants (squares denote median and error bars denote SEM). Furthermore, in the changepoint task, there is a higher learning rate from changepoint trials that decreases on the stable trials following a changepoint for both model (red) and human (blue) participants. Note how this qualitative behavioral trend is learned from the first observed changepoint (B) compared to the last observed changepoint (C). In contrast, in the oddball task shown in (D) the behavioral trend that is learned across the task is a decreased learning rate from the oddball trials compared to the following stable trials (lighter shades for the first oddball observed vs darker shades for the last oddball). (E,F) For the reversal task, we used a regression with update as the dependent variable and two independent variable: *Update*_*t*_ = *β*_0_ + *β*_1_(*Outcome*_*t*_ − *Prediction*_*t*_) + *β*_2_(*Mean*_*t*_ − *Prediction*_*t*_). For both model and human subjects, we see in (E) that with learning the coefficient for the first regressor decreases ( Model’s *β*_1_: early phase=0.39, late phase = 0.31,*p* < 0.0001, human participants *β*_1_: early phase=0.64, late phase = 0.57, *p* = 0.002) while in (F) the coefficient for the second regressor increases (Model’s *β*_2_: early=0.31, late =0.44, *p* < 0.0001, human participants *β*_2_: early=0.08, late =0.12, *p* = 0.19). (G,H) For the sequence learning task, we used a similar regression: *Update*_*t*_ = *β*_0_ + *β*_1_(*Outcome*_*t*_ − *Prediction*_*t*_) + *β*_2_(*Mean*_*t*_ − *Prediction*_*t*_). Here, again for both model and human subjects we see in (G) that with learning the coefficient for the first regressor decreases (Model’s *β*_1_: early phase=0.29, late phase=-0.02 *p* < 0.0001, human participants *β*_1_: early phase=0.42, late phase=0.23, *p* < 0.0001) while in (H) the coefficient for the second regressor increases (Model’s *β*_2_: early=0.17, late phase=0.75, *p* < 0.0001, human participants’ *β*_2_: early=0.21, late phase=0.38, *p* < 0.0001). The squares denote the group mean and the error bars denote SEM across simulations/participants.

### The behavioral signatures of structural learning

Given that both people and the pHDPi improved their performance with experience, we next sought to identify the specific characteristics of their learning behaviors that changed in order to afford that performance advantage. While more general measures of learning like sum of squared error show evidence of learning during a task, each task structure also has its unique characteristics of temporal structure learning. To look more closely into these unique signatures of learning, across model and participants, we used several behavioral analyses to look for temporal structure learning within each task.

We were able to find signatures of temporal structure learning across all four tasks: The main difference between changepoint and oddball tasks, in terms of the adaptive learning that they require, is the demand for a high (changepoint) versus low (oddball) learning rate in response to a surprisingly large prediction error. We observed that both the model and humans modify their learning rate dynamics differentially during learning changepoint and oddball tasks. To do so, we aggregated the data from all participants’ prediction errors and updates on the very first change point (or oddball) and last changepoint (or oddball) in the experiment and used this aggregated data matrix in separate regressions (early vs late) to measure trial update as a function of previous trial prediction error for several trials after the changepoint (oddball). Initially (Fig.5b,d, light colors), the trial-wise learning rate after the first observed state transition looks similar for changepoint and oddball tasks and is relatively flat indicating limited adjustments of learning in response to the state transition. However, for the last changepoint, the learning rate was highest on the trial after a changepoint (Fig.5c) whereas it was lowest for the trial after an oddball event (Fig.5d) consistent with the normative predictions for these tasks. Interestingly, the learning rate for the trial following the very first changepoint was elevated for people and the pHDPI (Fig.5b; learning rate for the first trial after first changepoint, human participants’ LR = 0.72, CI = [0.65, 0.78], model’s LR = 0.62, CI = [0.60, 0.65], learning rate for second trial after first changepoint human participants’ LR = 0.60, CI = [0.50, 0.70], model’s LR = 0.46, CI = [0.42, 0.50]). This initial predisposition toward changepoint-appropriate adaptive learning may reflect an inductive bias employed in the absence of structural knowledge, an idea supported in the pHDPi by minimal parameter changes required to accommodate changepoint structures. Overall, these results demonstrate that people and the pHDPi used experience within a structural environment to shape their reliance on surprising data points, emphasizing them in the changepoint structure and de-emphasizing them in the oddball structure, but also default to a learning policy closer to that demanded for changepoints.

Similarly, humans and the pHDPi model show evidence of temporal structure learning for the reversal and sequence learning task. For the reversal and sequence task, an agent that inferred the true structure should use knowledge from previously experienced latent states to predict the next outcome. In our task framework, this would correspond to relying on the mean of the relevant latent state to make predictions, rather than simply relying on recent prediction errors. We quantified this behavioral characteristic using a linear regression to explain updates after state transition in which one regressor was the difference between previous and current observations (prediction error) and the second regressor was the difference between previous prediction and mean of the current state (prediction to mean). If participants and the model learn to accurately predict the mean of the current state, we expect the prediction to mean regressor to have a higher coefficient toward the end of learning a task. Alternatively, if they relied solely on their preceding prediction error to adjust learning (i.e. no structure learning) then the prediction error coefficient should remain high. In the reversal task, the pHDPi showed a very clear shift from PE to inference-based updates (compare early versus late, red squares in Fig.5e,f; Prediction – outcome coefficient: early phase = 0.39, SEM = 0.01, late phase = 0.31, SEM = 0.01, *p* < 0.0001, Prediction – mean coefficient : early= 0.31, SEM = 0.02, late phase=0.44, SEM = 0.02, *p* < 0.0001). In the sequence learning task, the pHDPi shifted similarly from a prediction error based learning to direct inference (compare early versus late, red squares in Fig.5g,h; Prediction – outcome coefficient : early phase=0.29, SEM = 0.01, late phase=-0.02, SEM = 0.02, *p* < 0.0001, Prediction – mean coefficient : early=0.18, SEM = 0.01, late phase=0.75, SEM = 0.04, *p* < 0.0001).

The transition from relying on the *recent prediction error* to relying on the *mean* of the current state for updating the response, was also reflected in human participants’ behavior. Specifically, for human participants in the reversal learning task, the coefficient corresponding to prediction error effect decreased (compare early versus late, blue squares in Fig.5e; Prediction – outcome coefficient : early phase=0.64, SEM = 0.018; late phase = 0.57, SEM = 0.02, *p* = 0.002) while the coefficient corresponding to the difference between prediction and the mean on the reversal trials increased (compare early versus late, blue squares in Fig.5f; Mu – Prediction coefficient : early phase= 0.08, SEM = 0.02; late phase = 0.12, SEM = 0.03, *p* = 0.19). However, it is noteworthy that most participants, unlike the pHDPi, remained highly reliant on prediction errors in the reversal task even by the end of the experimental session – perhaps reflecting a bias toward this learning heuristic that deviates from normative structure learning (Tavoni et al., 2022). Human participants performing the sequence learning task also decreased their use of prediction errors (Prediction – outcome coefficient: early phase=0.42, late phase = 0.23, *p* < 0.0001; Fig.5g) and increased their direct inferences of the underlying mean (Prediction – mean coefficient: early=0.21, SEM = 0.02, late phase=0.39, SEM = 0.04, *p* < 0.0001; Fig.5h). Taken together, these analyses show that in tasks where latent states are recurring, participants learn to rely on the latent mean of these states, a hallmark of structural understanding.

### The pHDPi model explains asymmetric sequential effects in human structure learning

Recent experimental work revealed interesting and unexplained order effects on human learning observed after changes in the statistical structure of the environment (Bakst & McGuire, 2023). The study tested how people adjusted their gaze to predict the location of upcoming stimuli across two different structures (changepoints [CP], Oddball [OB]) that were experienced sequentially. Participants were divided into two groups: in one condition (CP→OB), the outcomes were generated from a Gaussian distribution with a fixed mean which occasionally was resampled from a uniform distribution. This condition was composed of two tasks in sequential order. The first task was similar to the changepoint task in our experiment) followed by a spontaneous un-signaled switch to an environment where the mean of the Gaussian was the outcome of the previous trial, i.e. a random walk process, with occasional outcomes being generated from a uniform distribution. Since this process is mathematically equivalent to the oddball task in our experiment, we will refer to it as the oddball task. The other group of participants experienced the same conditions, but in reverse order (OB→CP). They called this experience-driven modification in reacting to surprising events a “meta-learning” effect: People who switched from oddball to changepoint environments were able to increase their sensitivity to changepoints (higher learning rate for changepoint trials) while participants who switched from changepoint to oddball did not reduce their sensitivity to oddballs to anywhere near the same degree (Fig.6a-c).

**Figure 6.**
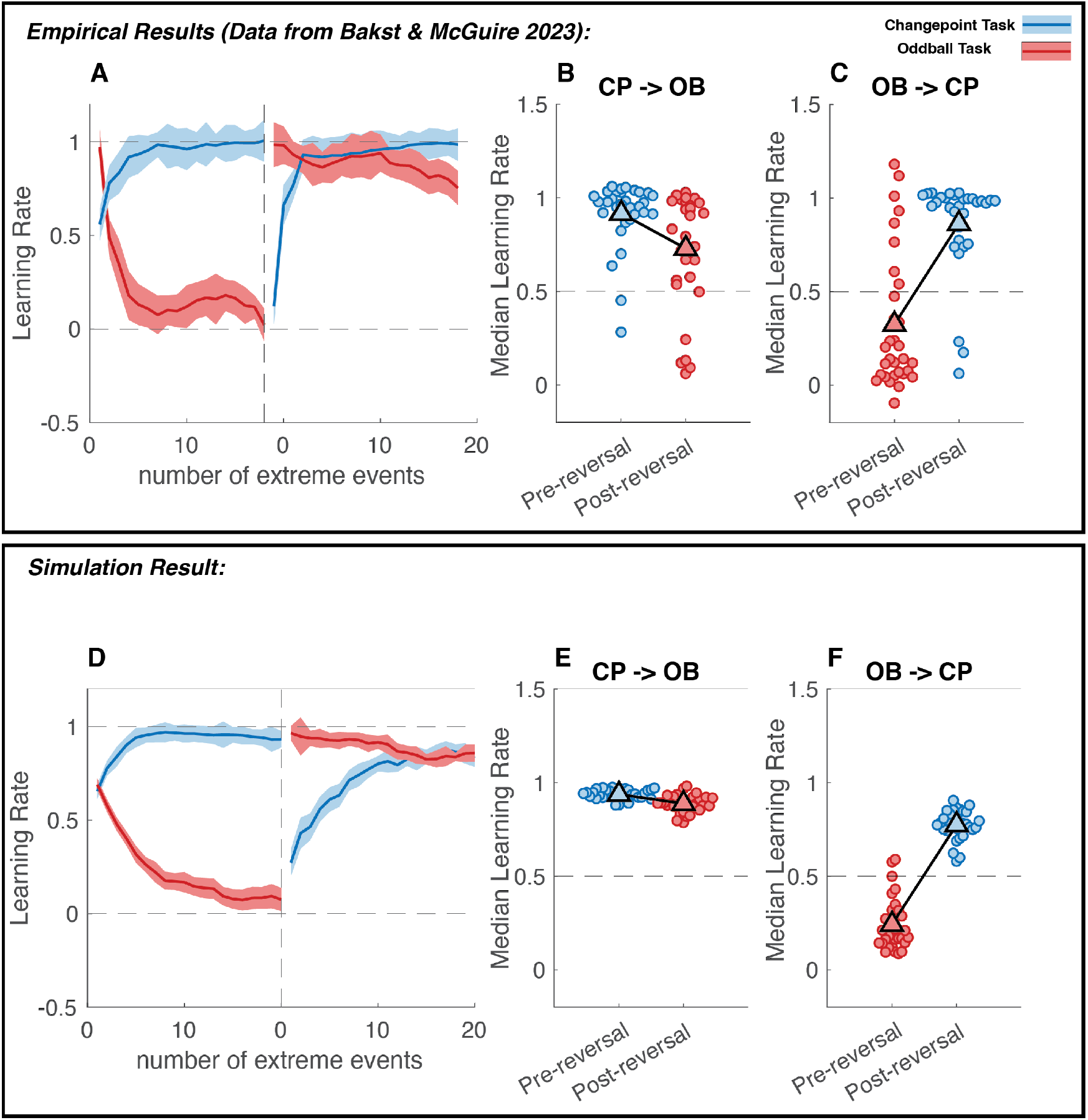
Humans and Model show order effects of task structure switch. A) Trial learning rate (ordinate) for extreme events (changepoints/oddballs) as a function of cumulative number of such events experienced (abscissa) for human subjects (data provided by Bakst & McGuire 2023 and used with permission). All participants experience changepoint (blue) and an oddball (red) structure but were divided into two groups that differed in the order of these conditions (transition depicted by vertical dashed black line). The task structures incentivized participants to use learning rates near one after changepoints and near zero after oddball events. Note that both groups of participants learned to use learning rates near one for the changepoint structure (blue), but those participants who experience changepoints first failed to converge on low learning rates in response to oddball events (red curve to right of dotted line). Solid line depicts smoothed group median and shading depicts SEM. B) The median learning rate across all extreme events is plotted for each human participant (circle) separately for changepoint (blue) and oddball (red) conditions and their order of presentation (B: CP → OB, C: OB → CP). Group mean values are depicted by triangles. Note the difference in median learning rate between the changepoint and oddball tasks (i.e. CP minus OB) is larger for participants who experienced the changepoint task first (CP →OB) in panel B than the group who experienced oddball task first (OB →CP) in panel C, (Wilcoxon rank sum p < 0.0001; average median LR difference in CP → OB = 0.19, average median LR difference in OB → CP = 0.54). D) Single trial learning rate (ordinate) for extreme events as a function of cumulative number of extreme events experienced (abscissa) for pHDPi model simulations of the same task setup. (E, F) Median learning rate for model simulations is depicted as circles and group mean as triangles and sorted according to condition and the order that conditions were encountered. Note the difference in median learning rates between conditions displays the same asymmetric learning effect: the difference is smaller in the CP→OB, panel E, than in the OB→CP condition, panel F (Wilcoxon rank sum p < 0.0001; average median LR difference in CP → OB = 0.05, average median LR difference in OB →CP = 0.53).

We reasoned that if the pHDPi model learned the statistical structure of changepoint and oddball tasks similar to humans, it would display asymmetric learning effects when transitioning from one task to another. In order to test this prediction and compare our model’s behavior with Bakst & McGuire findings, we tested the pHDPi model on a predictive inference task that included a structural transition. In this version, the model first saw 200 trials of a changepoint or an oddball environment, the generative structure of which follows Bakst & McGuire’s experiment, followed by 200 trials of the opposite (oddball or changepoint respectively). Similar to the human experimental set-up, the model was not provided with any cue with respect to the change in the generative structure. We simulated one run of the pHDPi model for each human participant (n=32 in each condition) and observed that the order of the tasks influenced how much the model adjusted its learning rate from extreme events in each task (Fig.6e, f). Specifically, we calculated ΔLR as per-subject median learning rate for CP minus their median learning rate for OB tasks (each median calculated across extreme events within that task structure). The ΔLR was smaller for the models which first experienced the changepoint and then the oddball tasks than the models which first experienced oddball, and then the changepoint tasks consistent with a failure to adapt to the new structure in that condition (Wilcoxon rank sum p < 0.0001, CP → OB: average ΔLR = 0.05, OB → CP: ΔLR = 0.53). The same analysis applied to human participants showed a similar result (Wilcoxon rank sum p < 0.0001, CP → OB: average ΔLR = 0.19, OB → CP: ΔLR = 0.54). These results suggest that pHDPi model formed inductive biases based on pretraining in a similar fashion to humans.

To identify the source of these inductive biases in the model we examined the hyperparameter inference in the pHDPi over the course of the experiment. We found that the pHDPi performing the CP → OB condition was unable to reduce its persistence sufficiently after the task switch (Fig.7c). Intuitively, this failure likely resulted from inherent ambiguity mapping model hyperparameters onto observed outcomes; the repeated presentation of the “typical” state in the oddball structure could either be interpreted as evidence for a single highly concentrated common state (low Gamma) or, alternatively, as evidence for a high level of state persistence. When the oddball task was learned individually, the pHDPi appropriately inferred low values of Gamma and Persistence (Fig.4d, e), however when it encountered the oddball task after pretraining on the changepoint task, the high inferred persistence levels were supported by the long strings of trials reflecting the “typical” state, thereby preventing the model from updating structural assumptions. To test whether the persistence hyperparameter was indeed the reason the model was unable to adapt post-switch in CP → OB condition, we repeated the same sequential task but manually updated the persistence (*η*) parameter to its oddball-appropriate value immediately after the CP → OB structural transition. This single change enabled the model to adapt its behavior appropriately post switch (Fig.7d-f). Taken together, these results demonstrate a dark side of structure learning observed in humans and explained by our model, namely that strong preconceptions about hyperparameters, when poorly calibrated to the current environment, can bias causal interpretations of new outcomes impairing predictions and learning.

**Figure 7.**
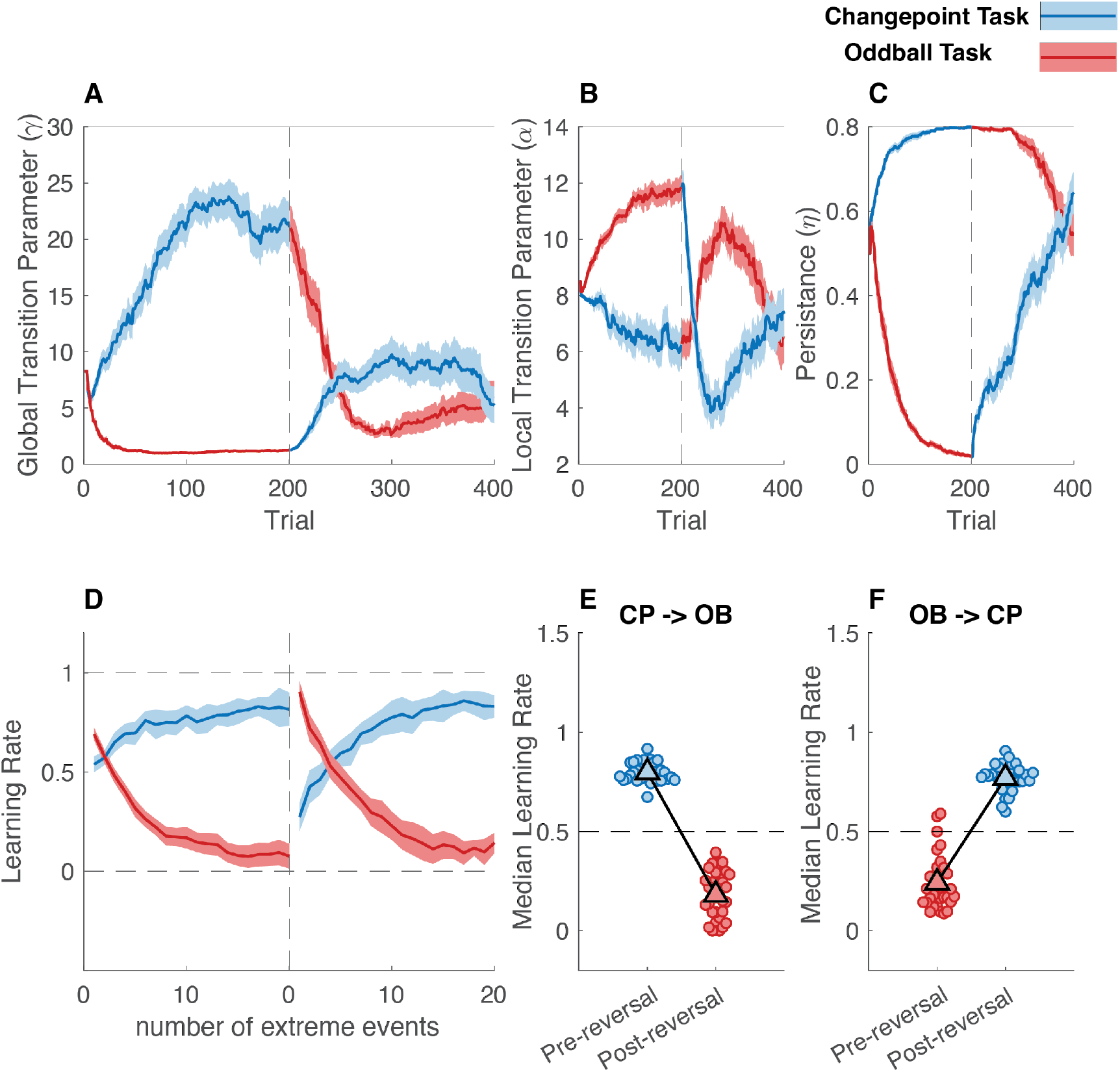
The persistence hyperparameter is responsible for the order effects observed in the model. A-C) Average values of inferred gamma (*γ*), alpha (*α*) and eta (*η*) in n=32 runs of model for CP → OB condition in blue, and OB→ CP condition in red with shaded regions denoting SEM and condition switch shown in dotted black line. D) Trial learning rate (ordinate) for extreme events as a function of cumulative number of extreme events experienced (abscissa) as in Fig.6d. Here we used the final inferred eta (*η*) value from pre-switch phase of OB→ CP condition in post-switch phase of CP → OB condition. Note how, unlike Fig.6d, the model adjusts its learning rate to the optimal value in OB task, post-switch. Solid line depicts smoothed group median and shading depicts SEM. e and f) Median learning rate for extreme events in simulated models, with single model’s learning rate depicted as circles and group mean as triangles. The differences in median learning rates are now similar in the modified model that experienced CP →OB condition in (e) and the model that experienced OB →CP condition in (f).

## Discussion

The world often undergoes changes that follow predictable temporal structures. We demonstrated that these structures can be learned and utilized to promote qualitatively different context appropriate adaptive learning behaviors. Specifically, we developed a normative model, the pHDPi, capable of solving structure learning through Bayesian inference over a compact but expressive prior for transition matrices (Fig.2). The model achieved high predictive accuracy across a range of contexts (Fig.3) by inferring hyperparameters most consistent with the observed outcomes, with key parameters pertaining to the total number of states, the similarity of transitions to states from each state, and the frequency of self-transition probabilities (Fig.4). Inference over hyperparameters allowed the model to gradually differentiate between changepoint, oddball, reversal and sequence structures – and to display context appropriate learning behaviors for each structure (Fig.5). These hyperparameters served as priors guiding inference in completely novel situations, such as when encountering a new state for the very first time.

Beyond a normative solution, the pHDPi provided a number of testable predictions for human structure learning. We used a predictive inference task framework to provide empirical evidence that people can use temporal structures without explicit information about environment structure labels. As predicted by the pHDPi, people leveraged experience to improve performance making predictions amid different temporal structures, with the smallest performance improvements encountered in changepoint tasks, which elicited the smallest hyperparameter adjustments in the pHDPi (Fig5.a). It is worth noting that the starting point of all the parameters was the expected value of the uniform prior over each parameter’s range, which would probabilistically define a truly “agnostic” model to any environmental structure. While this range was chosen somewhat arbitrarily for some parameters (e.g. gamma, alpha, noise or drift rate where in theory, there is no mathematical upper bound) we believe we chose an appropriate range for the types of tasks we considered ecologically relevant.

Furthermore, the performance improvements were accompanied by signatures of adaptive learning qualitatively matching those undergone by the pHDPi, albeit with additional biases, for example toward over-use of prediction error corrective learning (Fig.5b-h). We also found that in learning some structures (a sequence task), there’s a sudden phase transition from “unlearned” to “learned” behavior whereas in other tasks (a changepoint task), there are more gradual and less significant learning effects. The pHDPi not only provided insight into adaptive behaviors, but also into apparent learning failures. In particular, pretraining the pHDPi on one temporal structure and applying it to another led to a specific learning asymmetry observed in human behavior (Bakst & McGuire, 2023), namely an inability to deal appropriately with oddball events after training on an environment with changepoints (Fig.6). In this situation, the model revealed an inability to “unlearn” strong expectations for persistence, that caused the model to overreact to oddball events (Fig.7). Taken together, our results establish the first normative model of generalized structure learning and use it to elucidate the computational mechanisms through which people adjust their behavior in complex environments embedded with varied and unknown temporal structures.

Our work provides a meta-learning perspective that has often been ignored, or examined only for circumscribed cases, in previous work on adaptive learning (Bakst & McGuire, 2023; Behrens et al., 2007; A. Collins & Koechlin, 2012; d’Acremont & Bossaerts, 2016; Mathys, 2011, 2011; Nassar et al., 2010, 2019, 2021; Piray & Daw, 2020; Razmi & Nassar, 2022). A hallmark of adaptive learning is rationally responding to change. By addressing the problem of how a set of rational responses to change are learned de novo, we provide a potential solution to the “meta-learning challenge” of adaptive learning. A critical aspect of this solution is learning the right inductive biases over state transitions rather than learning the state transition matrix itself. While learning the state transition matrix or an approximation to it has been the focus of many recent studies aimed at understanding model-based behaviors (Gläscher et al., 2010; Momennejad, 2020), it is the inductive biases over the transition matrix, in our model controlled by inferred hyperparameters, that can be generalized to new states, or even new tasks, providing a sort of meta-learning. Here we show that such meta-learning can improve behavior when structures are unknown and stable by providing guidance on how to interpret a completely new state (Fig.4&5), but on the other hand can impair learning when meta-learned expectations are poorly matched to reality (Fig.6&7). Although we are not the first to point out this meta learning effect in human participants behavior (Bakst & McGuire, 2023; Simoens et al., 2025), we believe we are the first to provide a comprehensive account in terms of structure learning and a unified model that can deal with this heterogeneity of temporal structures. Taken together, our model and behavioral results provide a characterization of the meta-learning that enables people to learn to make predictions in qualitatively different environments.

Our results also have implications for the computational mechanisms governing adaptive learning. The pHDPi can describe adaptive learning behaviors across environments, and does so through a form of latent state inference. In this account, which has recently been suggested by several other lines of work, apparent adjustments of learning rate observed experimentally are attributed to the inference of an alternate context (Heald et al., 2020, 2023; Razmi & Nassar, 2022; L. Q. Yu et al., 2021), rather than to explicit adjustments of learning rate as had been proposed by a large body of previous work (Diederen et al., 2017; Mah et al., 2024; Nassar et al., 2010, 2012; Preuschoff & Bossaerts, 2007).This distinction matters the most when speculating about the neural basis of adaptive learning. While some previous work focused on finding a neuromodulatory signal in the brain corresponding to a dynamic learning rate (Diederen et al., 2017; Iglesias et al., 2013; Nassar et al., 2012; A. J. Yu & Dayan, 2005), contextual inference posits that learning dynamics could best be examined by characterizing latent state representations and their dynamics in the brain, which has also been an active area of research over the past decade (Nassar et al., 2018; Schuck et al., 2016; Schuck & Niv, 2019; Vaidya et al., 2021). It is noteworthy that, while latent state transitions contribute heavily to the learning behavior in the pHDPi, the model also learns global values for the volatility and stochasticity, which could be combined to control a baseline learning rate situations where the latent state remains stable (Piray & Daw, 2021). And thus, while the model supports the notion that learning dynamics are driven in part by discrete transitions in inferred latent states, it does not preclude the notion that there could be overall learning rate signal controlling sensitivity to new information in the absence of such transitions.

The contextual updating undergone in our model has direct implications for continual learning, an active area of research in both biological and artificial learning (Holton et al., 2025). Non-stationarity of the environment is often deemed an obstacle to be overcome in learning, with artificial neural networks suffering from catastrophic forgetting when they are trained in blocked (but not interleaved) conditions. In our work, we utilize specifically this ordered presentation of latent states (since an environment where states are presented randomly, by definition, doesn’t have any temporal structure) to guide learning. In this sense, our framework more closely resembles biological learning systems which are less susceptible to catastrophic forgetting when trained on blocked tasks structure and in some cases even benefit from it (Flesch et al., 2018; Heald et al., 2023; Lu et al., 2024; Russin et al., 2022). The mechanisms leveraging knowledge about state transitions in our model are complementary to similar work in artificial neural network models (Hummos, 2023; Nagabandi et al., 2019) that could potentially reduce the impacts of catastrophic forgetting in artificial neural networks, and at the same time, make them learn in ways that more closely resemble biological systems.

While our modeling and behavioral analysis provide insights into the basis for human structure learning, they do have a few limitations. We focused on providing a normative solution for learning temporal structures without an explicit supervised signal and testing its key qualitative predictions with respect to human behavior. While we saw qualitative features of normative learning across our participants, we also saw considerable individual variation in human behavior (Supplementary Fig.2), suggesting deviations from normative learning. Such deviations may reflect differences in priors of individuals from the flat prior over hyperparameters with which we equipped the normative model, however it is also possible that individuals differ in the degree to which they engage in structure learning at all, for example because the computations required are effortful (Bruckner et al., 2025; Glaze et al., 2018; Tavoni et al., 2022). Future work could take advantage of our model to distinguish between these possibilities through experiments where each participant experienced multiple structures, allowing for the distinction between a biased structure learner (e.g. one with a strong prior over hyperparameters) and one that makes no attempt to learn structure at all.

## Summary

Our results establish a generalized normative model for temporal structure learning and demonstrate its ability to explain patterns of learning in people confronted with qualitatively different environments. These results highlight the ability of people to exploit knowledge about the structure of their environment to improve local learning and prediction and reveal potential pitfalls of such structure learning when environments deviate from structural expectations. Taken together, these results provide a bridge to link a robust literature on how people and brains engage in adaptive learning behaviors appropriate for a given structure to the more open-ended problem encountered in real life, where structures themselves must be identified and learned de novo from experience.

## Acknowledgment

We thank Joe McGuire, Michael J Frank and members of Learning, memory and decision-making lab for helpful comments and feedback.

---

**Supplementary Figure 1.**
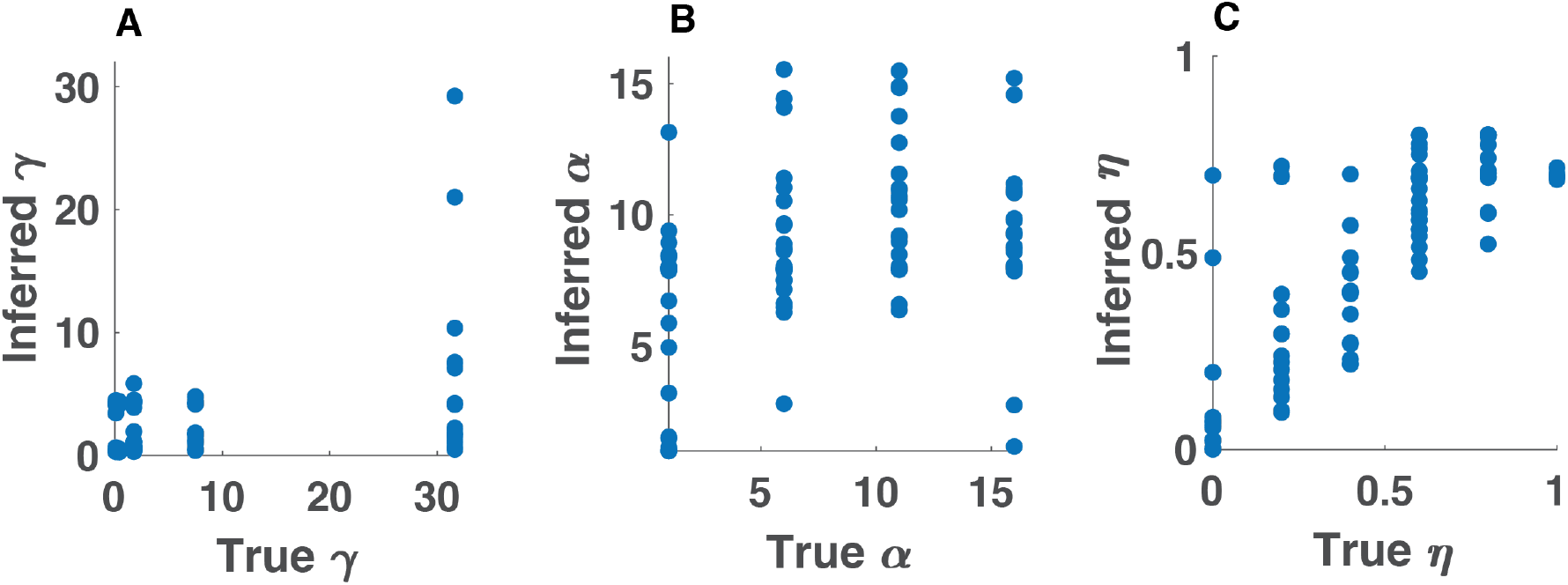
Parameter recovery for the pHDPi model. We randomly sampled 100 tasks from the ranges of hyperparameters used in the pHDPi model and generated 100 tasks with 400 outcomes directly from the generative process shown in Fig.2f using zero drift and low outcome noise (std = 10). We then ran the pHDPi model on generated tasks and plotted the averaged inferred parameter of the model (ordinate) versus the true hyperparameter of the generative process (abscissa). The model inferred parameter had a higher correlation for persistence (eta) and local transition probability (alpha) than global transition probability (gamma). The suboptimal recovery of gamma and alpha hyperparameters is to some extent expected since 1) the tasks are generated randomly, sometimes with no discernable structure, the empirical distribution of states might vary a lot and 2) by only using the outcomes, we are losing some of the information in the hierarchical generative process, where different combination of hyperparameters might be able to explain the results.

**Supplementary Figure 2.**
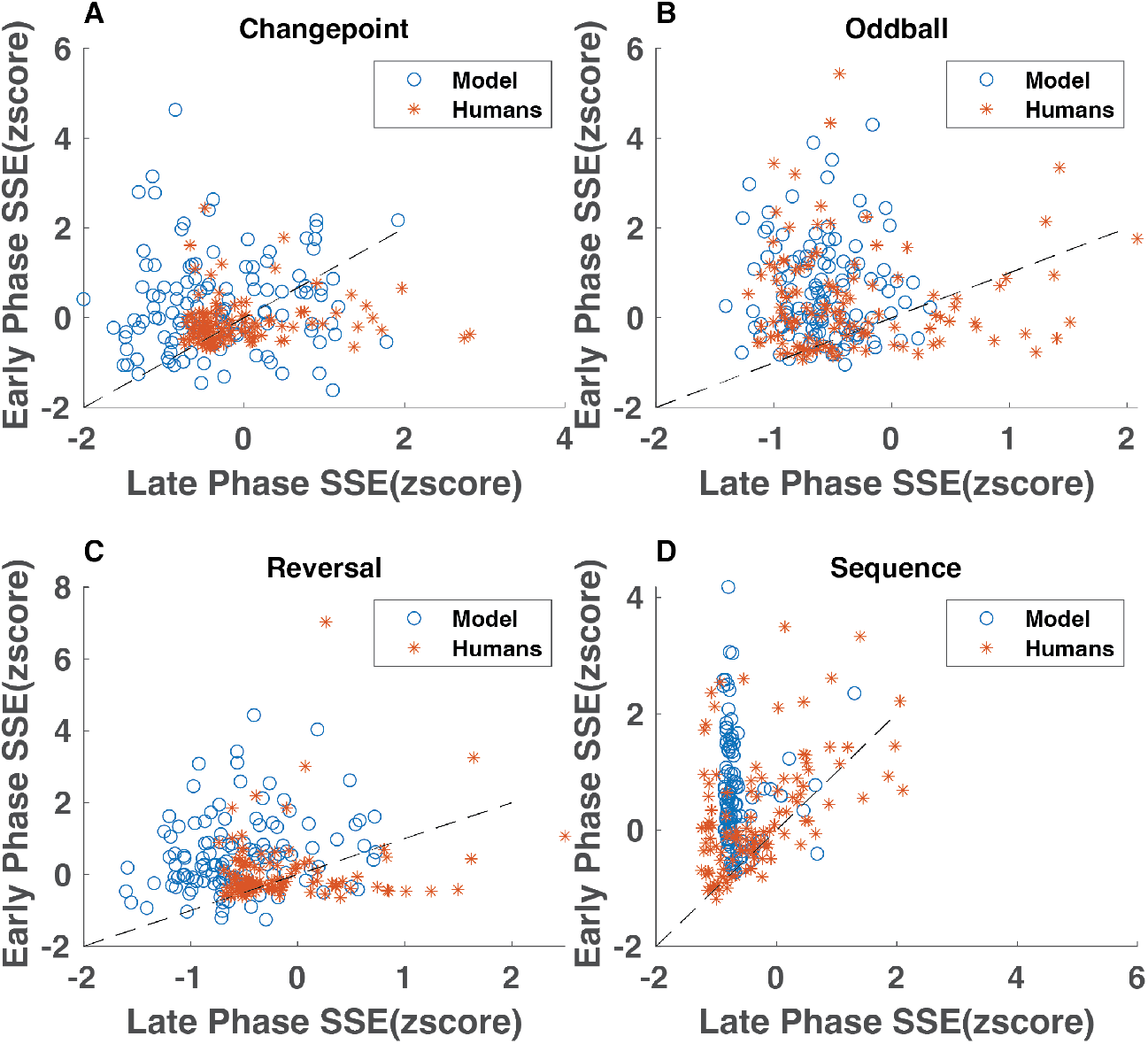
Individual variability in learning different tasks. Here, we plotted the same data shown in Fig.5a differently, with the z-scored sum of squared error over the first 30 trials (abscissa) versus z-scored sum of squared error over last 30 trials (ordinate) for: (a) changepoint, (b) oddball, (c) reversal and (d) sequence tasks, for human participants (red asterisks) and model (blue circles).While most human participants show improved performance (most red asterisks are above the identity line, i.e. larger early phase error), there is heterogeneity in how much humans learn in each task.

## Methods

### The structure learning task

We consider an online prediction task in which at each time point a random outcome *x*_*t*_ is generated from an underlying Gaussian distribution, and the task of the learner is to make prediction about next outcome, *p*(*X*_*t*_|*X*_1_, …, *X*_*t*−1_). This prediction problem would have been computationally straightforward if the distribution was constant. However, we consider the situation where the mean of the Gaussian distribution undergoes (different kinds of) change. For simplicity, we assume standard deviation is constant, however our model is flexible enough to deal with a changing standard deviation as well. We formalize “different kinds of change” by introducing transition probability matrices, where the rows of the matrix are probability of transitions from a state *Z*_*i*_ to any *Z*_*j*_ with Z denoting a categorical “state”. We assume that within each state *Z*_*t*_, all outcomes come from a truncated normal distribution (outcomes bounded by 0 and 100) with mean *u* and variance 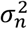.

Therefore, *X* is not IID distributed across the task but comes from a time varying distribution. For both model simulation and human behavioral experiments, we generated outcomes from a Gaussian distribution where the mean of each state was drawn from a uniform distribution 𝒰[0,100] and standard deviation was 25. The state transitions were according to the transition probability matrices shown in Fig.2b-e.

### Model

Our model is a Bayesian learner that starts with a prior probability over each row of the transition matrix, observes outcomes and uses Bayes rule to update that prior into a posterior belief. Inference over these hyperparameters allows the model to learn what state transitions to expect, even in cases where the model has never seen the specific state transition before.

#### Generative Process

The prior belief of the Bayesian model comes from a generative process. The true task of the modeler is choosing the right generative process. After choosing the right generative model, learning starts by inverting the prior with Bayes rule and doing inference. In this section we provide the detailed mathematical definition of the generative process (Fig.2f), introduced in the previous section specifying how states and their transition probabilities are generated as well as how outcomes are generated in a given state.

The persistent Hierarchical Dirichlet Process model assumes that observations in the environment are generated by a persistent Hierarchical Dirichlet process (pHDP). Described in more detail below, the pHDP model has a number of advantages for our overarching goals. First, it gives us a probability distribution over transitions to potentially infinite number of states, thereby making the model scalable. Second, it has sufficient flexibility to generate a wide range of qualitatively different transition matrices. Third, the expressiveness of the Hierarchical Dirichlet process is controlled by a relatively small set of parameters, each with an intuitive role in the generative process. In terms of inference, these properties suggest that inverting the pHDP should enable efficient inference over datasets of arbitrary size emerging from a wide range transition matrices by employing the appropriate inductive biases appropriate.

We draw responses from the Hierarchical Dirichlet process prior in three steps (Fig.2f):

1. First, we draw a probability vector *β* from a Dirichlet Process (DP) where ∑_*i*_ *β*_*i*_ = 1. The vector *β* is called the global transition probabilities and denotes the probability of each state in the task. We generate these probabilities by a stick-breaking process where we start with a unit length stick *β* and repeatedly break this stick by drawing a random variable *β*′_*k*_ from *Beta*(1, *γ*), multiplying the stick length by *β*′_*k*_, assigning the value to *β*_*k*_ and discarding this broken part of the stick. After *k* − 1 steps, the remaining stick length will be:

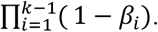

Higher values of gamma results in smaller broken stick portions and more of the original stick length remaining for subsequent time steps, therefore higher probability of new states in the future.
2. When a new state is created, the transition probabilities delineating the likelihood of visiting other possible states after the newly created state (i.e. the corresponding row in the transition probability matrix T) do not exist. One could think about creating these transition probabilities in a few different ways, and it’s useful to think about two extreme cases. On the one hand, we could a use the same vector of *β* that was used for the previously created state. In this case, the transition probabilities “out” of a newly encountered state will be exactly the same as those from all states, and thus we can infer them automatically. On the other hand, this generative process would be limited in its expressiveness – for example, it could never create a sequential transition matrix, where A→B, B→C, C→A. On the other extreme, we could draw a completely new vector of *β*s to describe transitions out of the newly encountered state. This generative process is more expressive, in that it allows for creation of a wider range of transition matrices, but it limits the degree of generalization possible – since transition probabilities from the newly encountered state would be conditionally independent of those in previously encountered states. Our approach can be thought of as a continuum between these extremes, as we introduce a parameter that allows us to generate outcomes ranging from highly expressive with limited opportunity for generalization on one extreme to minimally expressive with increased opportunity for generalization on the other. Specifically, we use the original *β* vector as the base of a second DP:

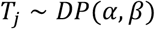

Where *α* is the concentration parameter and *β* is the base vector of the Dirichlet process. When the distribution is narrow (ie. *α* is large) transition probabilities from each state will be very similar to one another, affording efficient generalization, whereas when the distribution is wide (i.e. *α* is small) transition probabilities from each state will differ dramatically, limiting generalization but enhancing expressivity. The expectation of draws from this Dirichlet distribution will be the *β* vector. In order to generate rows of the transition probability matrix according to (1) we use a Chinese restaurant process where the probability of transitioning from each state i to previously visited state j is s:

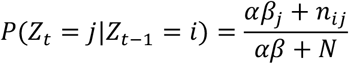

where *n*_*i*_*j* is the number of previous transitions from state i to j and *N* is the total number of transitions (i.e. trials) observed so far. If j is a new state, the probability of transitioning to it is given by:

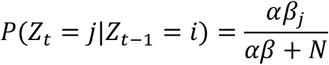
3. Lastly, to control the dynamics of self-transitions, we take a weighted average of the transition probability matrix obtained from steps 1 and 2 and the identity matrix of the same size (i.e. a matrix with ones on the diagonal and zeros everywhere else), where the weight of the second matrix is *η*. The third hyperparameter of the HDP, *η*, is a persistence factor that favors higher probabilities of self-transitions (the diagonal of the transition probability matrix).

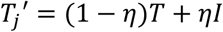

Where I denotes the identity matrix.

For each trial, the model makes a prediction about the mean of the outcome of current trial based on the probability of the active state from this prior, observe the outcome and update the prior by outcome likelihood, update the transition counts matrix by adding one to the number of observed outcomes in the active state, update the parameters within the active state (i.e. mean 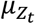) by outcome likelihood and drift the mean of the active state 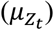 according to:

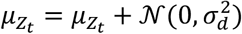

Where 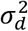 is the drift variance.

The hyperparameter *γ* or global transition probability parameter controls the overall probability of transitioning to a given state, where lower values of gamma results in one or a few states with high probability and higher gamma results in more probability of new states. In the second level DP, *α* or the local transition probability parameter controls how similar the rows of the transition probability matrix are to each other. With high *α* values, we get similar probabilities for transitions out of each state (so that where you will go next will not depend on where you are, this for example will not be helpful if you are in a sequence task). Finally, the hyperparameter *η* controls the degree to which the model tends to stay in one state in time irrespective of the popularity of that state.

#### Learning as inference

Given a task defined by its respective transition probability matrix and a Bayesian model that assumes the HDP generative process above, we define the problem of structure learning as the online inference of the five hyperparameters in the generative process. These include the three hyper parameters of the HDP i.e. global transition probability (*γ*), local transition probability (*α*) and persistence (*η*), the constant drift variance 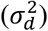 and constant noise variance 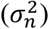 of outcomes.

That is we want to make a prediction of the current trial outcome by estimating the joint distribution of all the latent parameters of the task:

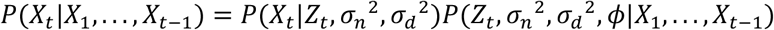

Where *X*_*i*_s are observed outcomes, *Z*_*i*_s are latent categorical states, *ϕ* is the three hyperparameters of the HDP: {*α, γ, η*} and 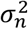 and 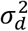 are noise and drift variance respectively. Each categorical state is defined by a probability distribution over the mean *u*_*z*_ and the emission distribution of each state is defined as:

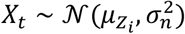

Specifically, we are interested in calculating the probability of a latent state given the outcomes observed so far using Bayes rule in a recursive manner:

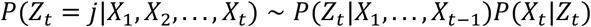

For making inference on the generative model explained in the previous section, we will use an online filtering approach for approximately estimating the posterior at each time step. Specifically, we start with t=1 and n equally weighted particles uniformly distributed on an equally spaced 5-dimensional grid of the free parameters of the model. Therefore, each particle is identified by a unique combination of free parameters 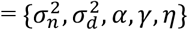. We will additionally copy each particle p times. These copies share the same free parameter ID and they are chosen to keep track of inference on the state sequence. This in total results in np particles. We then follow the steps below recursively and use the following approximation:

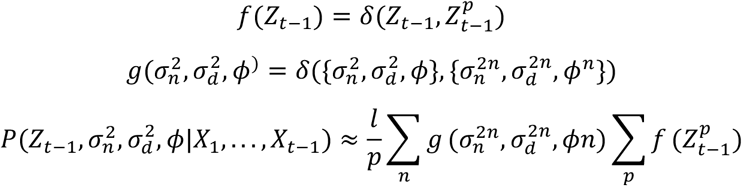

Where *δ* denotes the Kronecker delta function.

1 Propagate the particles one step forward in time according to:

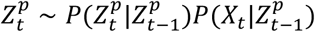

Where the second term on the right side is calculated as:

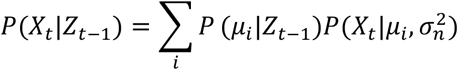

In the above equation, *u*_*i*_ is the discretized mean values. We omitted the particle *p* subscripts of all the terms for more clarity.
3 For particles that do not self-transition, update the transition probability matrix T by incrementing the corresponding *n*_*ij*_ of transition counts matrix and for particles that land on a new state update *β* by:
4 Within each group of particles with the same parameter ID, resample p particles with replacement based on their weights:

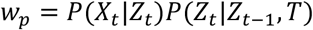

And setting the weights of the resulting particles to 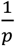 and *l*_*n*_ = ∑_*p*_ *w*_*p*_. The emission distribution (*X*_*t*_|*Z*_*t*_) is calculated similar to (9) and *P*(*Z*_*t*_|*Z*_*t*−1_, *T*) comes from the transition probability matrix.
5 Sample T and *β*:
  a. Sample the local transition probability matrix by:

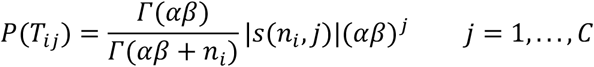

Equation (15) is the Chinese Restaurant Table (CRT) distribution, that is the PMF of tables in a Chinese restaurant process with *n*_*i*_ customers where s denotes Stirling numbers of the first kind and *n*_*i*_ is the total count of going from state i to other states.
  b. Sample global transition probability by:

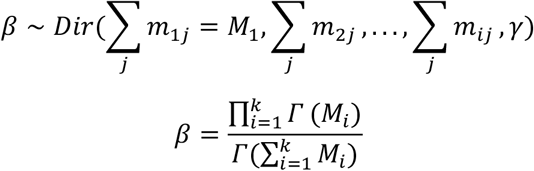
6 Go to 1, with t = t +1.

#### Alternative models

The delta rule models are defined by a constant learning rate :

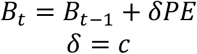

With *B* as the model’s estimate (or belief) and *PE* as the prediction error.

The Bayesian change-point model responds normatively in the changepoint task by using a dynamic learning rate:

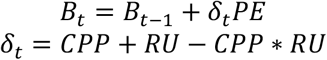

Here, *CPP* denotes the changepoint probability and *RU* denotes relative uncertainty. This model shows a learning rate of 1 on the changepoint trials, as *CPP* → 1 and a learning rate defined by the noise of the outcomes in the stable period as *CPP* → 0.

The Bayesian oddball model, responds normatively in the oddball task by using a different dynamic learning rate:

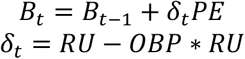

With *OBP* as oddball probability and *RU* as relative uncertainty. One the oddball trials, this model adopts a learning rate of 0 as *OBP* → 1 and otherwise maintains a learning rate proportional to the noise and drift of the outcome generating process.

#### Hyperparameter analysis

We used the model’s trial-by-trial expected value of each hyperparameter within a given experiment. For multidimensionality scaling (MDS) results, each hyperparameter was z-scored over all trials for each subject and adjusted by subtracting the first trial (starting point) value. This was to place all the different hyperparameters on the same scale for analyzing the temporal evolution of the model’s belief. We then ran MATLAB MDS function over this n by m matrix, with n = 480 for number of participants and m=5 for all the hyper parameters. For comparing the distance from the origin, we calculated the Euclidean distance for each task j, in 5 dimensional hyperparameter space as:

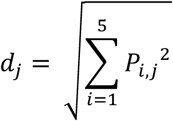

With *P*_*i,j*_ denoting the final z-scored and baseline centered estimate of hyperparameter *i* (= *γ, α, η, σ*_*N*_, *σ*_*d*_) for task *j* (= changepoint, oddball, reversal and sequence).

For calculating classification accuracy, we used MATLAB’s SVM function along with the matrix X containing all participants’ final hyperparameters. We ran four separate SVMs for each task, where Y was always binary vector defined as:

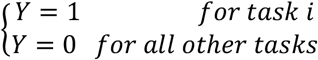

We reported the median accuracy of the SVM and repeated this procedure for each task 100 times to get bootstrapped SEM values.

#### Qualitative behavior analysis

Trial after changepoint learning rate regression: For the analysis presented in Fig.5b and c, we concatenated all the first (and last) changepoint trials across all participants (and model simulations) and looked at the trial wise learning rate, calculated as update over prediction error for the n-th trial after a changepoint.

For reversal and sequence learning tasks, we used a linear regression with update on one trial after state transition as dependent variable. We used two regressors as independent variables: 1) difference between previous and current observations and 2) the difference between previous prediction and mean of the current state:

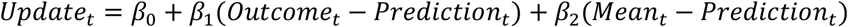

#### Sequential effects simulations

For the sequential effects results, we followed the experiment protocol of Bakst & McGuire 2023, second experiment, in which environment statistical structure reversed halfway through the experiment.

Briefly, in the original task, human participants were asked to report whether a briefly presented numerical digit was even or odd in order to obtain reward. Digits were presented on the periphery of a circle on the screen (These trials alternated with “central trials” in which digits were presented in the center of the circle. Since the original task included visual gaze analysis this was to ensure the participants maintained their gaze at the center in each trial. The gaze analysis is outside of the scope of the current paper and we refer readers to the original paper for more details about the experimental protocol.). The peripheral digits appeared in a Gaussian distributed location centered around a mean that was close the previous trial mean with high probability. Occasionally, the digits appeared on a random location anywhere on the circle. The generative process governing digit locations differed by condition: In changepoint (CP) condition, locations were generated from a Gaussian distribution with standard deviation 11.25 in visual angle, where the mean of the Gaussian was uniformly distributed from the circle perimeter with overall probability of 0.125. In the random walk condition, the mean was the previous digit location with outlier occurring anywhere on the circle with the same probability as in CP condition. i.e. 0.125. Participants were presented with three blocks of one condition followed up by three blocks of the other conditions in a counter balanced manner. Importantly, participants were not informed about task conditions, the structure of the blocks, or the halfway-reversal. Human data and analysis code was provided by the authors.

Trial-wise learning rate was calculated as:

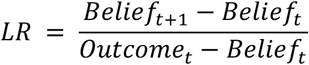

In the pre-reversal and post-reversal analysis, any trial-wise learning rate outside of the group inter quantile range by more than 1.5 IQR was discarded, separately for each phase. The group median was smoothed with a 5-trial window. Final learning rate was defined as median learning rate of the second half of the last block for both pre and post-reversal phase.

In order to simulate the same experiment with the pHDPi model, we created a task with 400 trials where the initial 200 trials were generated from a changepoint condition and the second 200 trials were generated from a generative process identical to Bakst & McGuire’s experiment. In our case, there were no blocks and no indication of condition change. The state means (helicopter locations) in our task were sampled from a uniform distribution U [0,300], and the standard deviation of the outcomes was 11.25. The overall probability of “extreme event” (changepoints/outliers) was similar to that in Bakst & McGuire’s experiment.

